# Polycomb represses a gene network controlling puberty via modulation of histone demethylase *Kdm6b* expression

**DOI:** 10.1101/2020.09.14.297135

**Authors:** Hollis Wright, Carlos F. Aylwin, Carlos A. Toro, Sergio R. Ojeda, Alejandro Lomniczi

## Abstract

Female puberty is subject to Polycomb Group (PcG)-dependent transcriptional repression. *Kiss1*, a puberty-activating gene, is a key target of this silencing mechanism. Using a gain-of-function approach and a systems biology strategy we now show that EED, an essential PcG component, acts in the arcuate nucleus of the hypothalamus to alter the functional organization of a gene network involved in the stimulatory control of puberty. A central node of this network is *Kdm6b*, which encodes an enzyme that erases the PcG-dependent histone modification H3K27me3. *Kiss1* is a first neighbor in the network; genes encoding glutamatergic receptors and potassium channels are second neighbors. By repressing *Kdm6b* expression, EED increases H3K27me3 abundance at these gene promoters, reducing gene expression throughout a gene network controlling puberty activation. These results indicate that *Kdm6b* repression is a basic mechanism used by PcG to modulate the biological output of puberty-activating gene networks.

## Introduction

A central event in the neuroendocrine cascade leading to the acquisition of reproductive maturity is an increase in pulsatile luteinizing hormone (LH) release from the pituitary gland (1). This increase is elicited by a change in the episodic discharge of gonadotropin hormone releasing hormone (GnRH), a decapeptide secreted by neurosecretory neurons located in the basal forebrain. In turn, GnRH release is driven by coordinated alterations in trans-synaptic and glial input to GnRH neurons (2, 3). A subset of neurons located in the arcuate nucleus (ARC) of the medial basal hypothalamus (MBH)(4–6) provides a primary trans-synaptic mechanism underlying pulsatile GnRH release. They are known as KNDy neurons, because they produce **K**isspeptin, **N**eurokinin B (NKB) and **D**ynorphin (4, 7).

Using a combination of DNA microarrays, genome-wide DNA methylation arrays, chromatin-immunoprecipitation (ChIP) assays, and gene expression analyses we recently discovered that an epigenetic mechanism of transcriptional repression operating in the ARC plays a significant role in timing female puberty (8). Our results identified the Polycomb group (PcG) of transcriptional silencers (9–11) as a major contributor to this repressive mechanism, and implicated EED, a member of the PcG repressive complex 2 (PRC2), as a core PcG component operating in KNDy neurons of the ARC to prevent the premature initiation of the pubertal process. Our results also showed that *Kiss1*, an essential component of the KNDy regulatory system, is a gene silenced by PcG, and demonstrated that this inhibition is lifted at the end of juvenile development allowing GnRH release to increase and puberty to take place (8).

To unveil the role of the PcG complex in the control of puberty we used a gain-of function approach aimed at perturbing the homeostatic make-up of the prepubertal ARC. While EED overexpression targeted to this brain region of the prepubertal hypothalamus demonstrated that *Kiss1* is a major PcG target, the gene networks that may be affected by EED, and that - expressed in either KNDy neurons and/or associated neuronal circuitries - contribute to the hypothalamic control of puberty have not been identified. In the present report we address this issue by using a systems biology approach. We first interrogated the MBH of prepubertal female rats overexpressing EED in the ARC using three different, but complementary approaches: massively parallel sequencing, high throughput targeted qPCR, and conventional RT-qPCR. We then analyzed the resulting data using computational methods able to identify and characterize genetic network architectures regulated by the PcG complex, without prior knowledge of the network(s) structure and function. Finally, we used chromatin immunoprecipitation assays and *in vitro* approaches to experimentally assess the validity of *in silico* predictions and gain insight into the biological significance of the changes in network structure caused by EED-driven perturbation.

## Results

### Physiological setup

The medial basal hypothalamic (MBH) tissue used for the present analysis was that previously reported (8). The MBH was dissected from 28-day-old female rats that had received six days earlier a bilateral intra-ARC microinjection of a lentiviral (LV) construct expressing rat *Eed* and Green Fluorescent Protein (GFP) under the control of the CMV promoter (LV-EED). Control animals received injections of the same LV expressing only GFP (LV-GFP). The ARCmedian eminence (ME) region of each animal was dissected and incubated for 4 h in Krebs-Ringer bicarbonate buffer at 37° under at atmosphere of 95% O_2_-5% CO_2_, with incubation medium collected every 7.5 min for measurement of episodic GnRH release (8). At the end of the incubation, the ARC-ME fragments were collected and processed for isolation of total RNA, which for the purpose of the present study was used for massively parallel sequencing (RNA-seq), quantitative high-throughput PCR (Open Array platform), and targeted RT-qPCR.

### Weighted Gene Co-Expression Network Analysis Reveals a Relationship Between *Eed*, *Kdm6b and Kiss1 Expression*

To discover potential co-expression modules of genes whose expression was altered by elevated levels of EED in the ARC, we subjected the RNA-seq data to Weighted Gene Co-Expression Network Analysis (WGCNA) (12). For this analysis we utilized the log2 counts per million (CPM) per sample of the top 5000 most variable genes having a minimum average CPM of 1 or >1, as summarized by the voom function (13) of the edgeR package (14) (http://www.R-project.org). The clustering pattern of these genes is illustrated in **Fig.1a**. We also examined alterations in expression pattern as illustrated by the eigengenes of each WGCNA-module (**Fig. 1b**), and utilized DAVID analysis (15, 16) to determine if genes contained in modules with altered expression in the ARC of *Eed*-overexpressing animals also showed functional overrepresentation.

**Figure 1.**
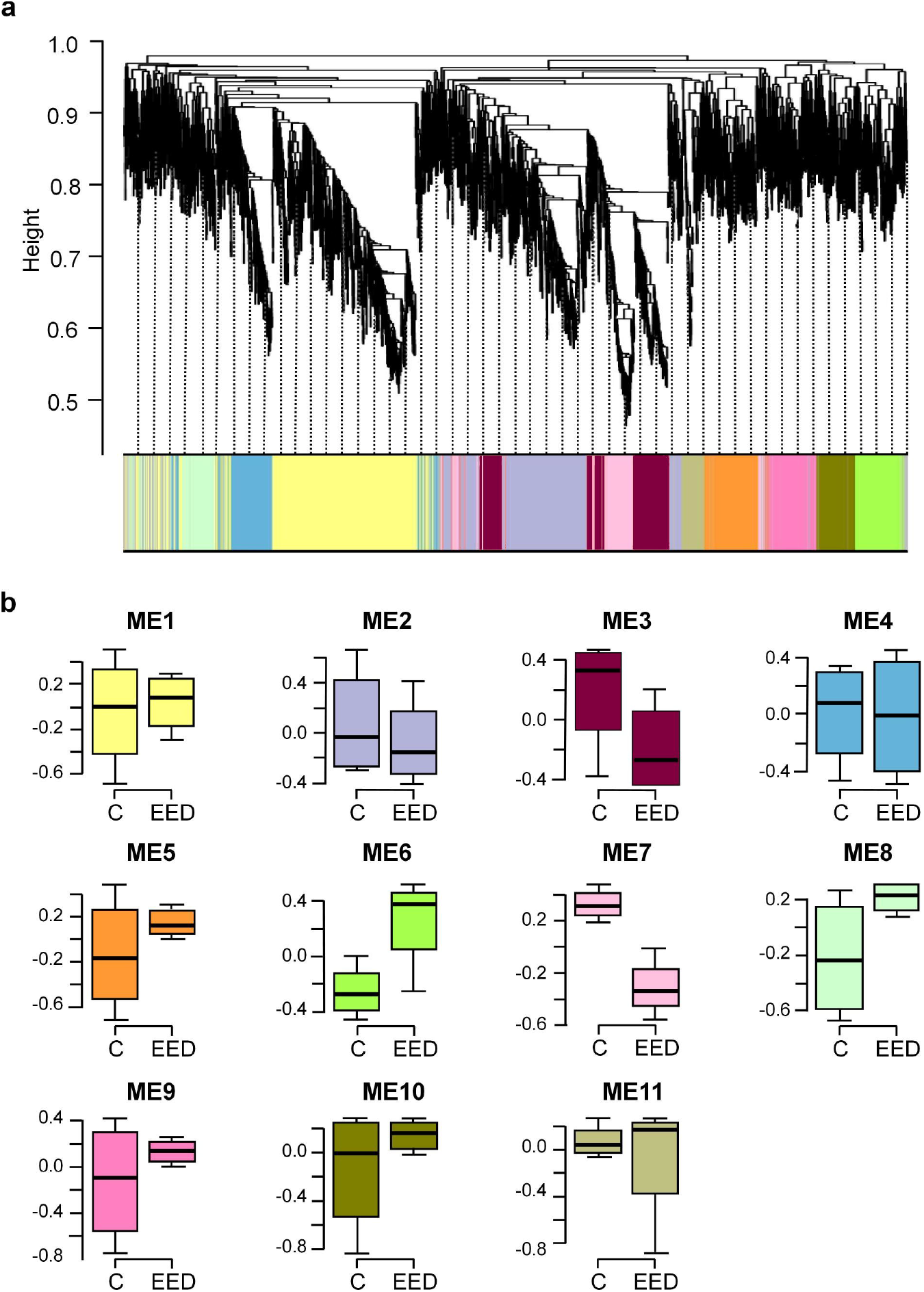
WGCNA analysis of gene expression after EED overexpression targeted to the ARC of prepubertal female rats. (**a**) Dendrogram of WGCNA module memberships of the top 5000 most variable genes detected by RNA-seq in the MBH of prepubertal female rats overexpressing EED in the ARC region of the MBH. Modules are identified by the colors depicted in 1b. (**b**) Boxplots of module eigengene values for control (C, LV-GFP-injected animals) and EED-overexpressing (EED, LV-EED-injected animals) groups (excluding module zero). Boxplot colors map to the dendrogram shown in 1a.

Only modules 6 and 7 exhibited clear separation of overall expression between LV-EED and LV-GFP-injected animals, with module 6 upregulated and module 7 downregulated in the MBH of rats injected with LV-EED. Functional overrepresentation analysis indicated that module 6 was strongly enriched for terms related to signaling, immune response and myelination pathways, while module 7 showed relatively little functional enrichment (**Supplementary Table 1a, b**). However, several individual members of these co-expression modules were genes that could be important for the epigenetic regulation of pubertal timing. For instance, module 6 (which, as expected, contains *Eed*), also includes *Kat2b*, a gene that encodes an acetyltransferase shown to stabilize the PRC2 complex of PcG (17) and *Mbd4*, which encodes a protein that recognizes and binds methylated DNA (18). Module 7, on the other hand, contains *Kiss1* and *Kdm6b* (**Supplementary Table *2***), an intriguing co-localization because the major function of KDM6B is the demethylation of H3K27me3 at promoter regions targeted by the PcG complex (19, 20). From a mechanistic perspective, we found the hypothesis that KDM6B could regulate *Kiss1* expression by acting as an antagonist of PcG-dependent gene silencing compelling in light of our previous results showing that EED directly represses *Kiss1* activity in KNDy neurons of the ARC (8). The changes in expression for these and several other genes observed after EED overexpression were all nominally significant (**Supplementary Table 2**).

### EED decreases Glutamatergic Gene Expression

We noted several genes encoding glutamatergic receptors that were down-regulated by EED overexpression (**Supplementary Table 2**), including clusters of these genes associated with WGCNA modules 2 and 3. This suggested that an important outcome of enhanced PcG repression is inhibition of glutamatergic transmission. To further understand the potential interplay of *Eed* and *Kdm6b* with genes encoding glutamatergic receptors, in addition to other genes of interest, we used an OpenArray platform to assay a targeted set of 224 genes with potential involvement in the regulation of puberty (21). In addition, we employed targeted qPCR to assess the expression of genes that were not represented on the OpenArray set or that were not accurately assayed in this platform due to either low levels of expression or imprecise replication.

This study demonstrated significant differential expression of several genes in the MBH of *Eed*-overexpressing animals as compared with the MBH of controls injected with LV-GFP (**Supplementary Table 3**). In agreement with the RNA-seq results, *Kdm6b* was downregulated in *Eed*-overexpressing animals. In addition, several genes encoding either glutamatergic receptors or molecules involved in glutamatergic transmission were heavily downregulated (**Supplementary Table 3**). Notably, *Nell2*, a gene selectively expressed in glutamatergic neurons and encoding a glycoprotein that promotes neuronal growth and supports glutamatergic signaling (22) was the single most downregulated gene among those assayed using the combination of Open Arrays and qPCR.

### *Kdm6b* Expression Is Related to Increased GnRH Pulse Frequency

To determine if *Kdm6b* expression is related to GnRH pulsatile release, we first analyzed the basic characteristics of pulsatile GnRH release from incubated ARC-ME fragments and observed that the frequency of GnRH pulsatility correlated strongly with total GnRH release (**Fig. 2a**), but not with the average amplitude of pulses. We then used a partial correlation analysis strategy to assess the existence of potential relationships between gene expression and GnRH secretion, and found that expression of several genes had a strong partial correlation with either total GnRH release or pulse amplitude when the influence of the other variable was removed (**Fig. 2b, Supplementary Table 4**). Of these genes, only *Kdm6b* showed a strong positive partial correlation with GnRH release after removal of pulse amplitude correlation as a variable, and a strong negative partial correlation with pulse amplitude, once the effect of correlation with total GnRH release was removed (**Fig. 2b**). Thus, higher *Kdm6b* expression appears to correlate with more frequent GnRH pulses, consistent with the strong correlation of pulse frequency with overall release noted earlier. This inference was supported by regression analysis; while *Kdm6b* expression on its own was a relatively poor predictor of total GnRH release, addition of a *Kdm6b* expression-pulse amplitude interaction term considerably improved the fit of the regression (**Fig. 2c**). An additional *Kdm6b* expression-pulse frequency term led to an even tighter fit to total GnRH release (**Fig. 2d**). Overall, these results suggest that *Kdm6b* plays a significant role in the regulation of GnRH release, possibly via positive control of *Kiss1* expression as suggested by our RNA-seq and Open Array results. However, it is also clear that regulation of *Kiss1* expression alone might not fully account for the alterations in GnRH release we observed. In particular, the partial correlations of *Kdm6b* with GnRH pulse frequency suggested that *Kdm6b* might be involved in regulating additional neuronal excitatory systems controlling GnRH release. The loss of glutamatergic receptor gene expression revealed by our RNA-seq and Open Array analyses (**Supplementary Table 2 and 3**) supports this assumption.

**Figure 2.**
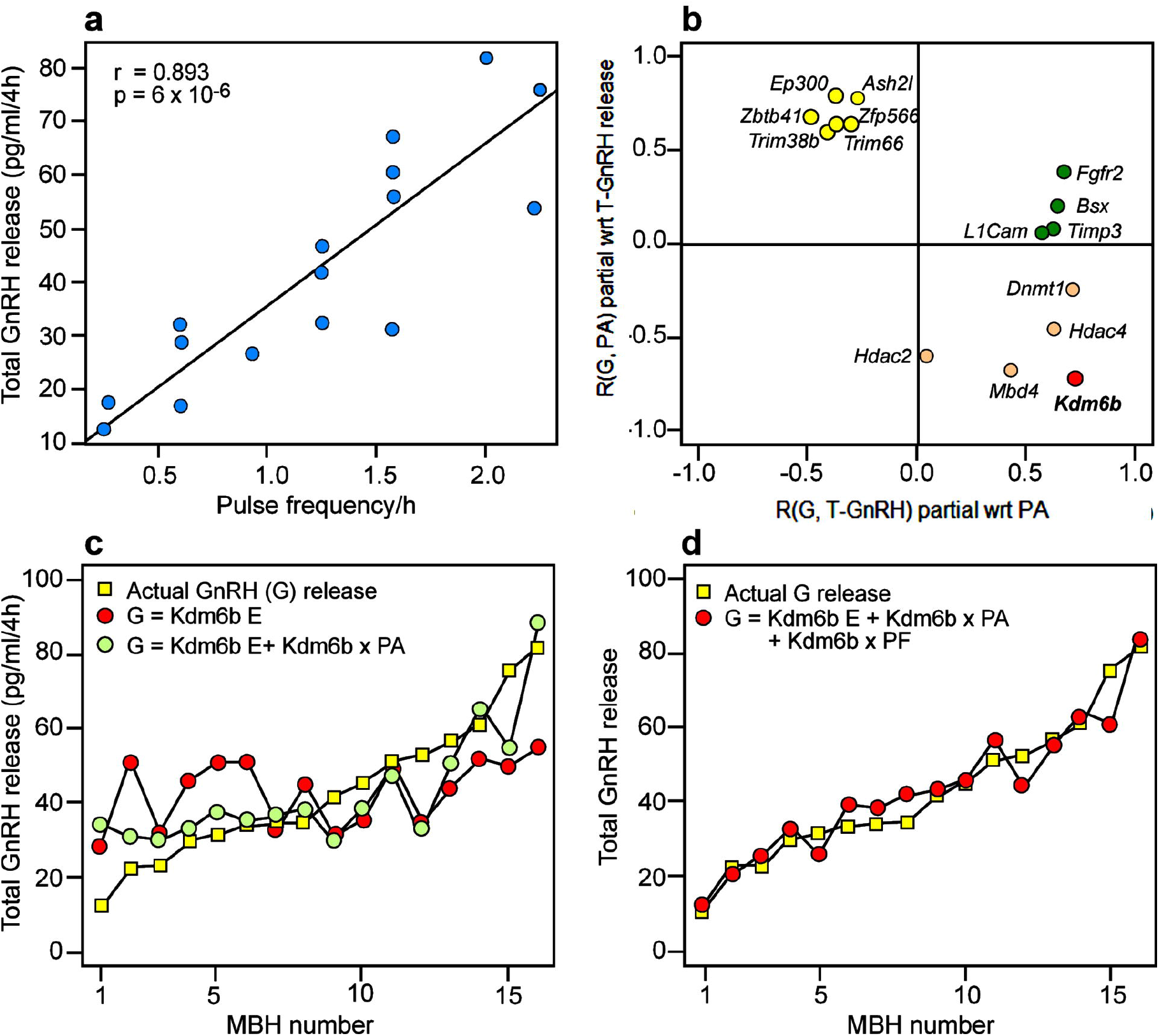
Correlations between gene expression and GnRH release from the MBH after EED overexpression. (**a**) Scatterplot of total GnRH release vs. average GnRH pulse frequency/hour over a 4 hour incubation period of MBH fragments derived from late juvenile 28-day-old female rats injected six days earlier with a lentiviral expressing GFP alone or EED plus GFP. Best-fit linear correlation is indicated by black line. (**b**) Plot of partial correlations (R) of gene expression (G) as assayed by OpenArray/qPCR with total (T) GnRH release and pulse amplitude (PA) with regard to (wrt) other physiological metric. (**c**) Plot of total GnRH release ordered from lowest to highest (yellow squares) and regression model predictions for Y□_Total Expression_ = β_intercept_ + β_*Kdm6b* expression_ (red circles) and Y□_Total Expression_ = β_intercept_ + β_*Kdm6b* expression_ + β_*Kdm6b* expression x Pulse Amplitude (PA)_ (green circles). (**d**) Plot of total GnRH release ordered from lowest to highest (yellow squares) and regression model prediction for Y□_Total Expression_ = β_intercept_ + β_*Kdm6b* expression_ + β_*Kdm6b* expression x Pulse Amplitude(PA)_ + β_*Kdm6b* expression x Pulse Amplitude (PA) x Pulse Frequency(PF)_ (red circles).

### PcG/ KDM6B are linked to *Kiss1/* Glutamatergic/Potassium Channel Gene Expression

In addition to the above described expression analysis, we performed a compressive sensing-based co-expression network inference on the OpenArray data and targeted RT-PCR data as described earlier (21). We utilized this method instead of WGCNA due to the smaller number of genes examined by our PCR analyses and the ability of the compressive sensingbased method to detect robust relationships between individual genes. The results of this analysis indicate a strong direct positive relationship between *Kiss1* and *Kdm6b* expression (**Fig. 3a**) that recapitulates the clustering of these two genes in module 7 of our WGCNA analysis. Importantly, both genes also showed a robust negative correlation with *Ezh2*, the catalytic member of the PRC2 H3K27-methyltransferase complex (9), supporting the hypothesis of an antagonistic role for KDM6B and the PcG complex (22) in the regulation of *Kiss1* expression. In addition, *Kiss1* showed a negative relationship with *Gatad1* (previously shown to repress puberty (21)), *Setdb1*, which encodes an H3K9-methyltransferase that catalyzes the synthesis of H3K9me3, a repressive histone mark (23, 24), and *Gabrag2*, the gene encoding the gamma2 subunit of a GABA_A_ receptor. This last observation is interesting, because alterations in GABAergic/glutamatergic signaling balance could be a factor underlying the relationship of pulse frequency and overall GnRH release we observed in our physiological experiments. Intrigued by this possibility, we searched for the second neighbors of both *Kiss1* and *Kdm6b* in the co-expression network to identify additional genes of interest.

**Figure 3.**
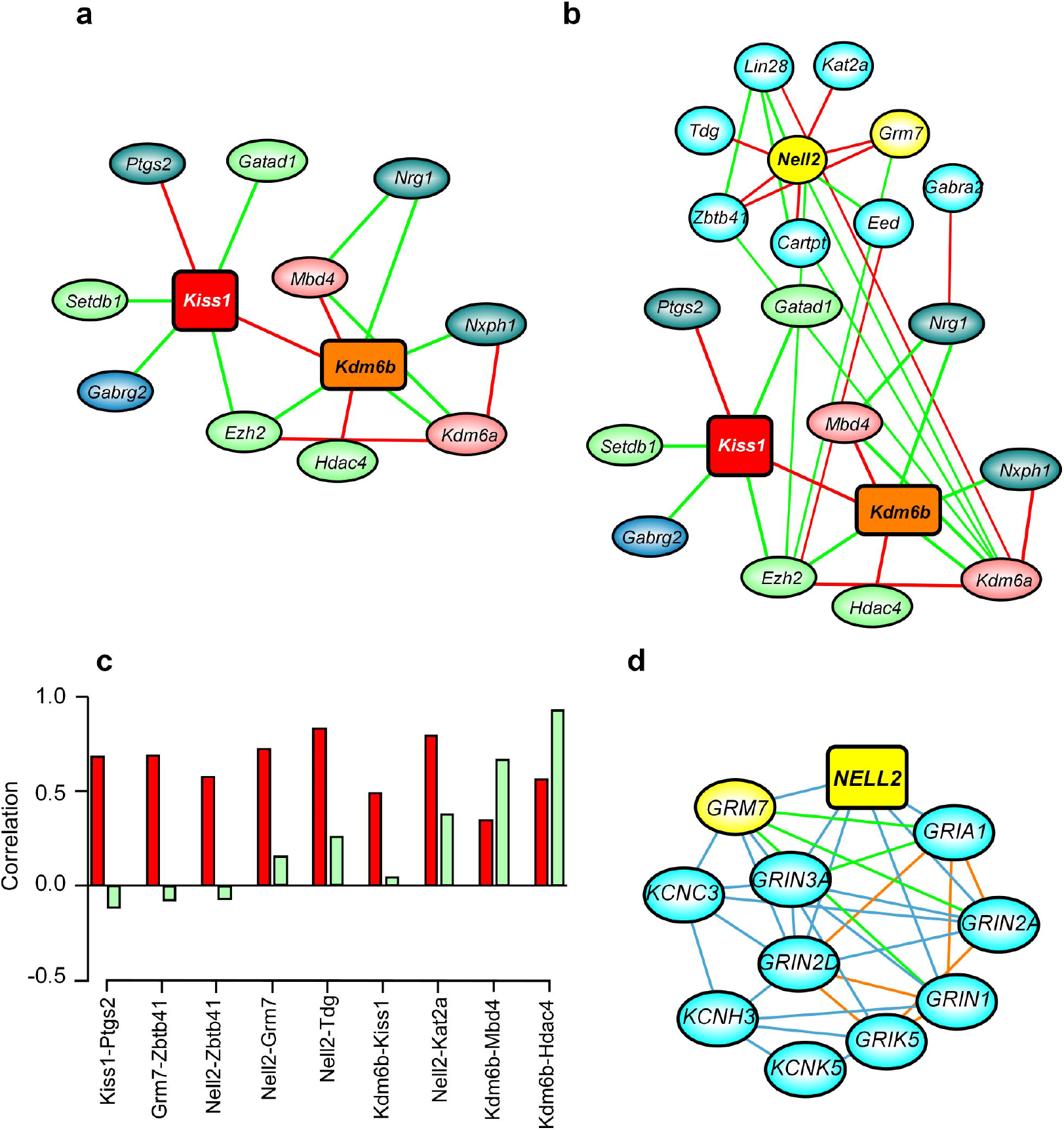
Gene co-expression networks in the MBH of immature female rats injected with either LV-GFP or LV-EED in the ARC. (**a**) First-neighbor network of strong co-expression edges for *Kiss1* and *Kdm6b*. Positively correlated edges are red, negatively correlated edges are green. First neighbors involved in primarily negative epigenomic regulation are indicated in light green ovals, while positive epigenomic regulators are indicated in red ovals. Non-epigenetic genes are blue ovals. (**b**) Second-neighbor network of strong co-expression edges for *Kiss1* and *Kdm6b*. Glutamatergic-related genes are indicated in yellow ovals, other in cyan ovals. (**c**) Histogram depicting expression correlations of pairs of genes involved in positively correlated strong co-expression relationships with *Kdm6b, Kiss1, Nell2* and *Grm7* under control conditions (red bars) and after EED overexpression (light green bars). (**d**) Network of previously-reported relationships between the glutamatergic-related genes *Nell2* and *Grm7* (yellow nodes) with glutamatergic and potassium channel genes downregulated by EED-overexpression (cyan nodes), detected by either RNA-seq or OpenArray, and shown as reported by the GeneMANIA database query tool. Blue lines indicate co-expression, green lines indicate genetic interactions and orange lines indicate known pathway interactions.

We found that another GABA receptor gene, *Gabra2*, is a second neighbor of *Kdm6b* and that two genes encoding proteins involved in glutamatergic transmission, *Nell2* and *Grm7*, are second neighbors to both *Kdm6b* and *Kiss1* (**Fig. 3b**). *Grm7* encodes a Group III glutamatergic metabotropic receptor involved in the etiology of mood disorders (25). Interestingly, *Nell2* and *Grm7* are direct neighbors of *Ezh2*, suggesting the likelihood of a regulatory relationship between *Kdm6b*, the PRC2 complex, and genes encoding glutamatergic receptors. Further analysis of these associations revealed that the majority of positively correlated co-expression edges between the highly-connected *Kdm6b, Kiss1, Nell2 and Grm7* nodes and neighboring genes observed in the MBH of LV-GFP injected animals diminished considerably in strength after EED overexpression (**Fig. 3c**). The exceptions were the positive correlations between *Kdm6b* and *Mbd4/Hdac4* expression as the strength of these associations was not reduced by EED overexpression. Altogether, these results suggest that the PcG complex keeps puberty in check not only by repressing *Kiss1* expression, but also by disrupting co-expression of functionally associated genes that interact with *Kiss1* within the boundaries of a gene network involved in excitatory neurotransmission.

In light of these findings, we re-examined the co-expression modules from our RNA-seq experiment and discovered that a number of glutamatergic genes showing at least nominally significant downregulation under EED-overexpression are components of modules 2 and 3 (**Supplementary Table 1c, d**). In addition, we noted a highly significant enrichment for potassium channel genes in these modules (**Supplementary Table 1c, d**). Strikingly, some of the potassium channel genes that were downregulated have been characterized as leak channels involved in maintaining neuronal membrane potentials near action potential thresholds (e.g *Kcnk4, Kcnk5*) (26) or with speeding neuronal recovery after action potentials through delayed rectification (e.g, *Kcna1*)(27). Additionally, *Kcnn1*, a calcium-responsive potassium channel involved in suppression of membrane excitability and regulation of spike train intervals (28) is located in the upregulated module 6, consistent with its oppositional role in regulating membrane potential compared to the majority of potassium channel genes located in downregulated modules.

Because many of these genes were not present in the OpenArray design we could not directly assess the existence of relationships with *Kdm6b* or members of the PcG complex. To overcome this limitation, we used the GeneMANIA database of known relationships between genes (29) and found strong co-expression, genetic interaction and pathway interconnection between glutamatergic and potassium channel genes downregulated by EED overexpression, including *Nell2* and *Grm7* (**Fig. 3d**). The concordance of this finding with the basic structure of our inferred strong co-expression network and WGCNA modules suggest that the genes most prominently co-regulated with *Kiss1* by EED/KDM6B belong to a cohort of genes involved in excitatory neurotransmission.

### Targeted RT-PCR Confirms Network Differential Co-Expression Predictions

To confirm the EED-induced changes in expression predicted by both WCGNA analysis of RNA-seq data and our compressive sensing-based co-expression network inference we used a targeted RT-PCR approach. In addition to *Kdm6b, Kiss1* (module 7, Fig. 1) and *Kat2b* (module 6, Fig. 1), we measured a subset of mRNAs encoding the glutamatergic receptors and potassium channels depicted in Fig. 3d. We compared the expression profiles of these genes in the MBH of control and EED-overexpressing animals *in vivo* with profiles observed in control R22 hypothalamic cells and R22 cells overexpressing EED *in vitro*. With exception of *Kcnn1* whose mRNA levels increased under EED overexpression *in vivo*, but strikingly decreased in EED overexpressing cells *in vitro*, the changes of expression induced by EED were similar *in vivo* and *in vitro* for all other genes analyzed (**Fig.4**). Surprisingly, *Nell2* and *Grm7* are strongly expressed in the MBH, but not in R22 cells (**Supplementary Fig.1**). These results indicate that genes encoding a defined subset of glutamatergic receptors and potassium channels are under PcG repressive control both in the MBH *in vivo* and in hypothalamic cells *in vitro*.

**Figure 4.**
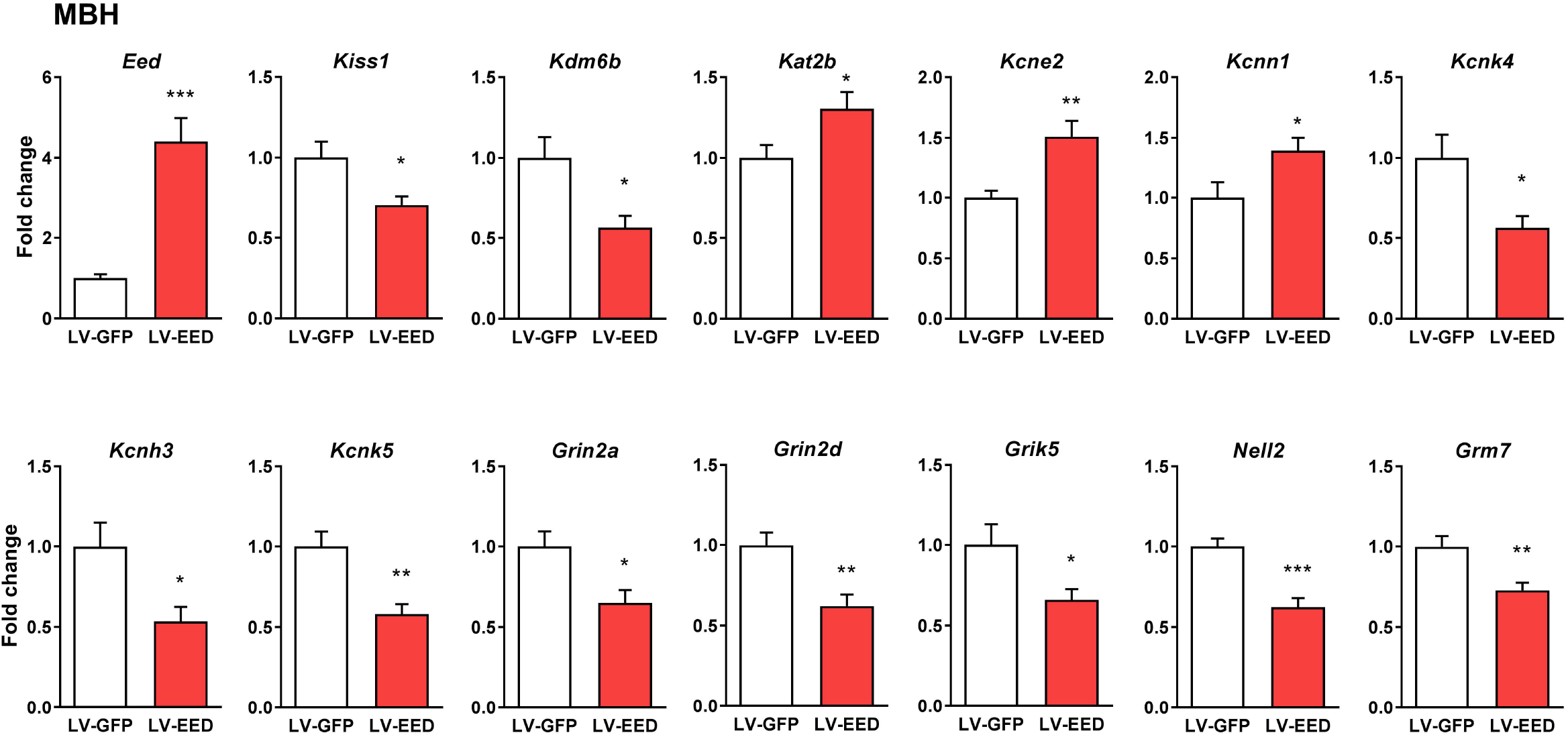
Changes in gene expression elicited by EED overexpression targeted to the ARC of immature female rats. Loss of expression of selected epigenetic, glutamatergic-related and potassium channel genes classified in modules 2, 3, 6 and 7 by WGCNA analysis, after overexpressing EED in the ARC of immature female rats. The animals received a bilateral injection of LV-GFP or LV-EED in the ARC at the beginning of juvenile development (22-days of age), and the MBH was collected on postnatal day 28; mRNA levels were measured by qPCR. Results are expressed as fold change with respect to control values. * = p < 0.05, ** p < 0.01, *** p < 0.001 vs. LV-GFP-Control treated rats. (Student’s t-test) (n=8 per group).

### EED increases H3K27me3 abundance at the promoters of network genes

While our physiological and co-expression analyses strongly indicate that EED/PRC2 represses the activity of genes important for pubertal development, they do not inform us as to whether or not EED is directly recruited to the regulatory regions of these genes. To address this question, we used a chromatin immunoprecipitation approach, utilizing R22 cells stably overexpressing EED.

We first determined if EED is recruited to the promoter regions of *Kdm6b, Kiss1* and glutamatergic and potassium channel genes previously identified as being EED targets (**Fig. 3d**). We then assessed H3K27me3 levels at the promoters, as a proxy for *Kdm6b’s* effect. Recruitment of EED to all the promoters examined increased significantly after EED overexpression (**Fig. 5a**). The content of H3K27me3 also increased (**Fig. 5b**), indicating that □ consistent with its role in PRC2 function (9) □ EED facilitates the deposition of H3K27me3 at the promoter region of downstream target genes. Overall, these results support the notion that the PRC2 complex down-regulates the promoters of several glutamatergic-related and potassium channel genes, in a manner consistent with its previously demonstrated role as a negative modulator of pubertal timing (8). Notably, the majority of the potassium channel genes repressed by EED have been shown to be involved in maintaining membrane potential or facilitating recovery after the action potential (26–28). This suggests that their regulation, along with regulation of glutamatergic genes, represents a physiological mechanism underlying the antagonistic relationship of *Eed* and *Kdm6b* expression with the loss of GnRH pulsatility that occurs in the presence of elevated EED levels in the ARC of animals approaching puberty.

**Figure 5.**
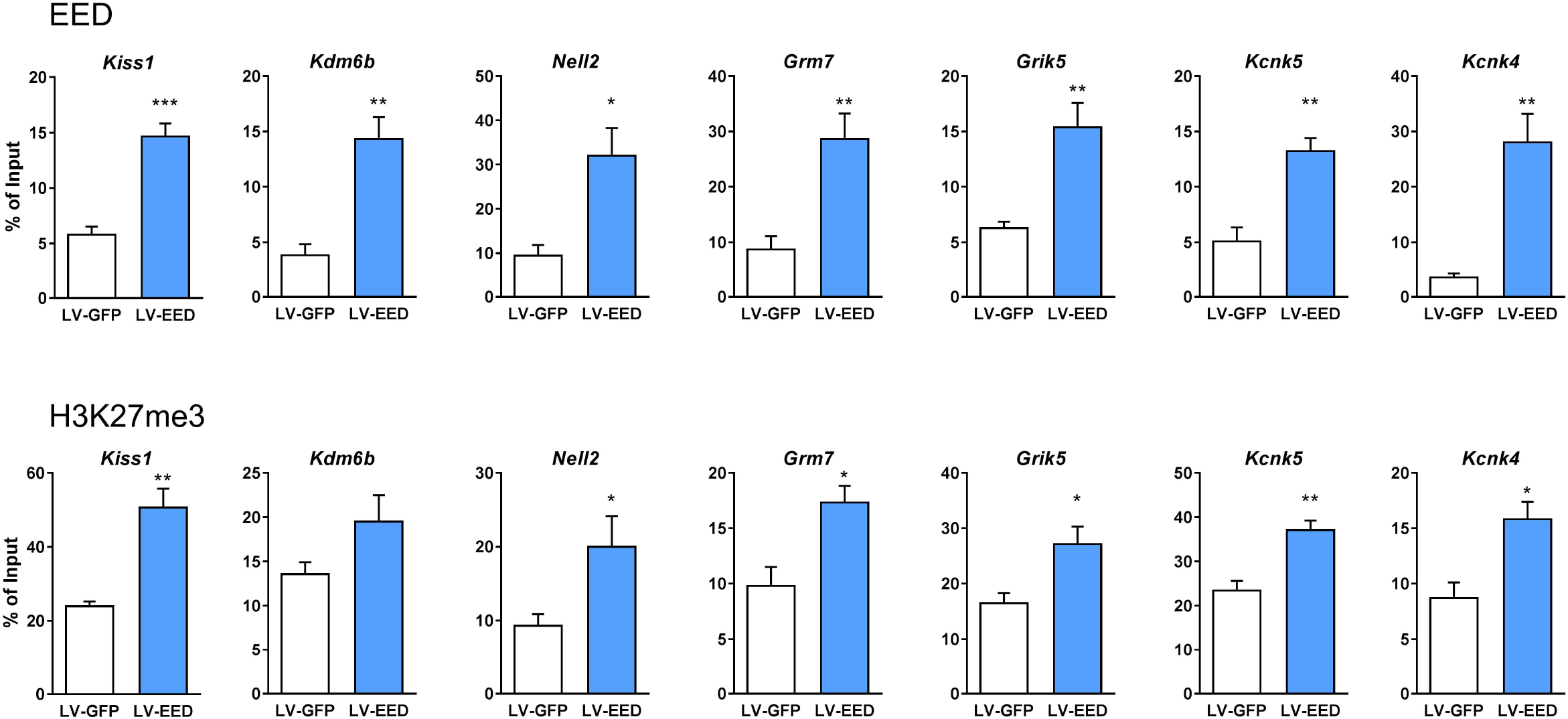
Recruitment of EED and H3K27me3 to the promoter of EED targeted network genes. (**a**) Recruitment of EED to the promoters of *Kiss1, Kdm6b*, glutamatergic-related and potassium channel genes showing decreased expression in R22 cells overexpressing EED. (**b**) Increased H3K27me3 abundance at the promoter of *Kiss1, Kdm6b*, glutamatergic-related and potassium channel genes that recruit EED in R22 cells after EED overexpression. Results are expressed as fold-change with respect to cells transduced with LV-GFP. * = p < 0.05, ** = p < 0.01, *** = p < 0.001 vs. LV-GFP treated cells (Student’s t-test) (n=4 per group).

### KDM6B counteracts EED-mediated effects on selected gene network members

From the above-mentioned *in vivo* and *in vitro* results, it became clear that by interacting with promoter regions and increasing H3K27me3, EED downregulates the expression of *Kdm6b, Kiss1* and other second tier genes involved in glutamate signaling and potassium dependent membrane transporters. To determine if these second-tier genes are directly affected by EED or by the loss in *Kdm6b* expression, we assessed the ability of *Kdm6b* to counteract the repressive activity of EED on a selected group of genes.

We observed that transient transfection of either *Eed* or *Kdm6b* into R22 cells leads to the expected increase in expression of these genes, as determined by RT-qPCR (**Fig. 6a and b**). We also observed that *Kdm6b* antagonizes the repressive effect of *Eed* on *Kiss1* mRNA expression (**Fig. 6c**), and partially, but significantly, counteracted the effect of *Eed* on *Grik5, Kcnk4* and *Kcnk5* (**Fig. 6d-f**). As indicated earlier these three genes are involved in excitatory neurotransmission and are co-regulated with *Kiss1* by EED/KDM6B (**Fig. 3**).

**Figure 6.**
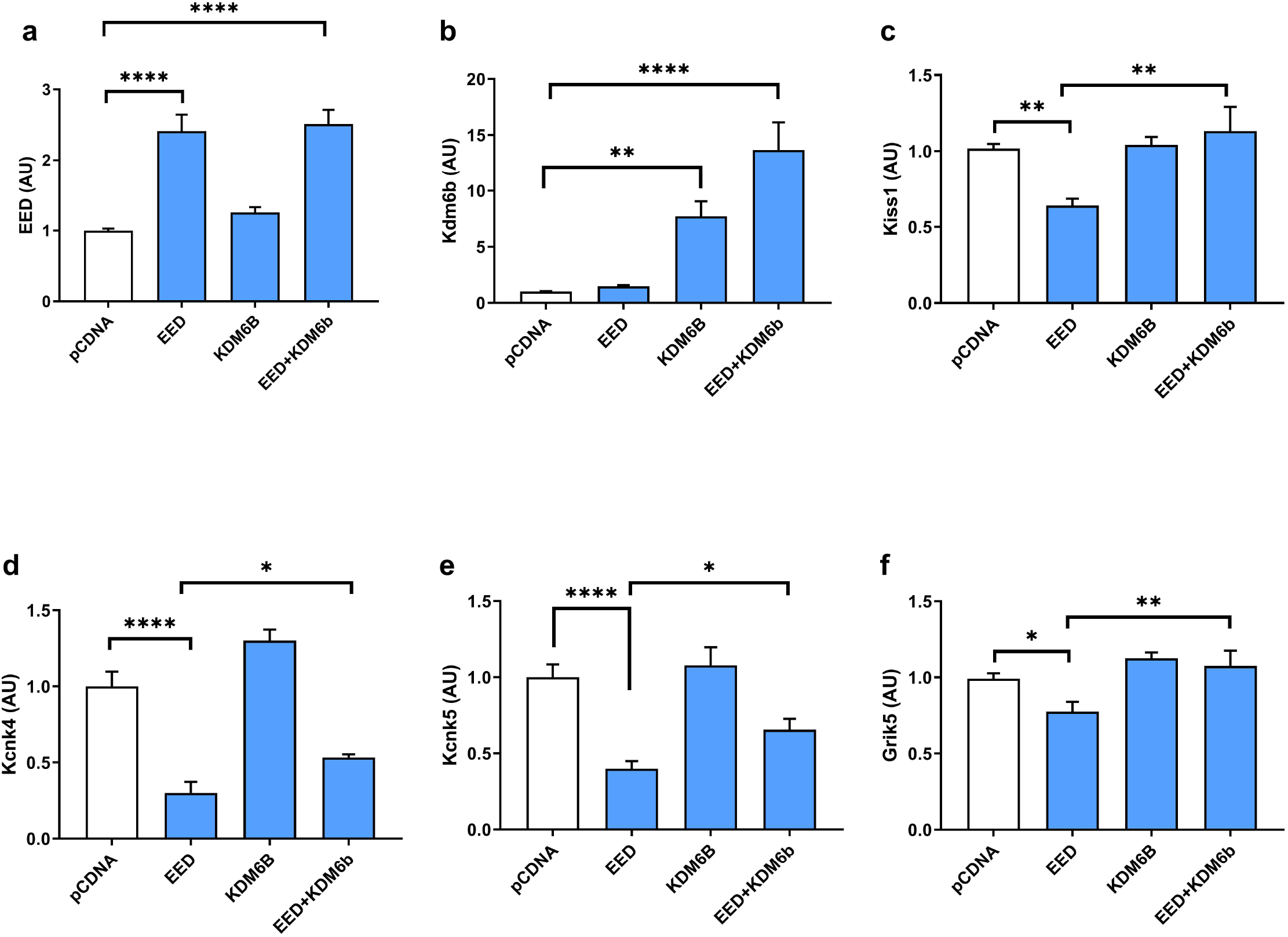
KDM6B counteracts the effects of EED. Forty-eight hours after transient transfection of Eed and Kdm6b expression vectors into hypothalamic R22 cells, mRNA levels were measured by qPCR. Results are expressed as fold change with respect to control values. * = p < 0.05, ** p < 0.01, **** p < 0.0001 vs. LV-CG-Control treated cells (Student-Newman-Keuls test) (n=3 per group).

Altogether, these results lend credence to the concept that the Polycomb complex keeps puberty in check by repressing not only the *Kiss1* gene, but also the expression of genes involved in the neuroexcitatory control of puberty.

## Discussion

In mammals, pulsatile GnRH secretion is a mode of neurosecretion characterized by the periodic release (every 30 min) of discrete amounts of GnRH into the portal circulation connecting the hypothalamus to the pituitary gland (30, 31). Increased GnRH pulse frequency is crucial for female reproductive function as it is required for both pubertal maturation and the follicular and preovulatory phases of the menstrual cycle in adults (32, 33). These episodes of GnRH release closely follow bursts of multi-unit electrical activity in the ARC (31, 34–36), and require the coordinated action of the neuropeptides neurokinin B/kisspeptin and dynorphin (37, 38) to occur. Although some of the genetic networks coordinating GnRH neuronal activity have been identified (3, 39), little is known about the epigenetic mechanisms that may coordinate gene networks involved in facilitating GnRH release during pubertal development. The present report addresses this issue.

We have previously demonstrated that a Polycomb group-dependent epigenetic mechanism of transcriptional repression operates within the ARC nucleus of the hypothalamus to time the initiation of female puberty. This repressive influence leads to diminished pulsatile GnRH release, and involves transcriptional inhibition of the *Kiss1* gene, which is crucial for GnRH release (8). Peripubertal female rats overexpressing *Eed (a* component of the Polycomb Repressive Complex 2) in the ARC showed decreased *Kiss1* expression, increased GnRH inter-pulse interval, decreased total GnRH release and delayed puberty (8). Thus, when EED abundance in the ARC increases GnRH secretion decreases and puberty is delayed.

We now provide insights into the integrative mechanisms underlying the epigenetic regulation of puberty by identifying a role of KDM6B, a histone demethylating enzyme, in the control of pulsatile GnRH release before the acquisition of reproductive maturity. The Polycomb complex catalyzes trimethylation of histone 3 at lysine 27 (H3K27me3), and use this histone modification to repress gene expression (19, 40). In turn, KDM6B regulates this process by erasing the K27 methylation mark from H3, and thus reducing the prevalence of H3K27me3 at gene regulatory regions (20).

Using a gain of function and a system biology approach we discovered the existence of a genetic network in the ARC that displays *Kdm6b* as a central node, with *Kiss1* and several epigenetic related genes (*Ezh2, Hdac4, Kdm6a, Mbd4*) as first neighbors. Our results also demonstrated that *Kdm6b* not only regulates the expression of *Kiss1*, but also the transcriptional activity of a cohort of genes involved in excitatory neurotransmission, and therefore in the stimulatory neural control of GnRH release (39–41). These genes encode glutamatergic receptors (*Grm7, Nell2, Grik5, Grin2a/d, Grin1* and *Gria1*), a glutamate release-inducing molecule (*Nell2*), and potassium channels (*Kcnh3, Kcnc3, Kcnk4/5*) responsive to arachidonic acid metabolites (42) and present in the hypothalamus (43), where they facilitate neuronal excitability.

A remarkable feature of this KDM6B-dependent regulatory system is the strong correlation that exists between *Kdm6b* expression and GnRH pulse frequency, as revealed by the correlation analysis of 5000 genes changing expression in the ARC after exposure to elevated levels of EED. Functional analysis of *Eed/Kdm6B* interactions revealed that EED inhibits gene expression by recruiting the repressive H3K27me3 histone mark to gene promoters expressed in the ARC. When *Kdm6b* content is enhanced, trimetlylation of H3 K27 is diminished and the repressive effect of EED on *Kiss1* and the network genes involved in excitatory neurotransmission is lost. These results are, therefore, consistent with the view that *Kdm6b* functions in a highly dynamic manner to facilitate the rhythmic activity of GnRH neurons during pubertal development, and controls GnRH pulse frequency by antagonizing the repressive effect of the PcG complex on genes involved in the stimulatory control of GnRH secretion.

Previous reports have highlighted the importance of an EED/KDM6b regulatory system in the control of two other critical development events, the first mammalian cell lineage commitment (41), and mammalian embryo implantation (42). By demonstrating the involvement of the EED/KDM6b system in the control of GnRH output at puberty, our findings provide an additional example of a critical developmental event in the lifespan of an individual subject to tight epigenetic control.

## Methods

### Animals

The animals used were those described in a previous publication (8). They were juvenile (22-28 days of age) Sprague Dawley female rats, and were obtained from Charles River Laboratories international, Inc. (Hollister, CA). Upon arrival, the rats were housed in a room with controlled photoperiod (12/12 h light/dark cycle) and temperature (23–25°C), with *ad libitum* access to tap water and pelleted rat chow. The use of rats was approved by the ONPRC Animal Care and Use Committee in accordance with the NIH guidelines for the use of animals in research.

### Preparation of EED-expressing lentiviral particles

A detailed description of the procedure we used to prepare lentiviral particles expressing the rat *Eed* gene (LV-EED) was reported earlier (8).

### Stereotaxic delivery of LV-EED

LV particles carrying an *Eed* transgene were stereotaxically delivered to the arcuate nucleus (ARC) of the hypothalamus of 22-day-old female rats as reported (8).

### Tissue collection

The ARC-ME fragments used for incubation were dissected as reported (8), i.e. by a rostral cut along the posterior border of the optic chiasm, a caudal cut immediately in front of the mammillary bodies, and two lateral cuts half-way between the medial eminence and the hypothalamic sulci. The thickness of the tissue fragment was about 2 mm.

### GnRH release and gene expression in ARC-median eminence (ME) explants in vitro

For this analysis, we used previously reported GnRH values derived from ARC-ME fragments of 28-day-old female rats that had received LV-EED into the ARC on PND22, and had been incubated *in vitro* for 4h to assess pulsatile GnRH release in samples collected every 7.5 min (8). We used these ARC-ME fragments to extract total RNA for qPCR analysis of gene expression.

### Correlation analysis of GnRH release and Gene Expression

We used a partial correlation approach using the ppcor package in R to determine partial correlations of gene expression as determined by OpenArray with total GnRH release and pulse amplitude after removing the correlation of the other physiological output. The ppcor test function was used to evaluate the significance of each partial correlation, and the significance values were then adjusted using the Benjamini-Hochberg multiple testing correction.

We also performed a regression analysis on the relationship of Kdm6b expression to total GnRH output using a standard linear regression model. Models included the intercept and a Kdm6b expression term, as well as either an interaction term for Kdm6b expression and GnRH pulse amplitude or both Kdm6b/pulse amplitude and Kdm6b/pulse frequency interaction terms.

### EED overexpression in R22 hypothalamic cells

#### Lentivirus infection in vitro

The ability of the LV-EED to alter gene expression and trigger changes in abundance of H3K27me3 at putative target promoters was examined using the immortalized R22 hypothalamic cell line (Cedarlane, Burlington, NC). The cells were plated in DMEM medium at 400,000 cells/well using 12-well plates. Twenty-four hours later, the cells were transduced with the viruses at a multiplicity of infection (MOI) of 5 to 1. Control cells were transfected with LV particles expressing enhanced green fluorescence protein (eGFP) under the control of the CMV promoter and lacking Eed (LV-GFP). Three days after the infection, transduced cells (identified by their expression of eGFP) were isolated by flow cytometry to produce a pure population of cells. These cells were expanded and re-plated onto 12 well plates at a density of 300,000 cells/plate. Three days later, the cells were collected, aliquoted and stored at −80°C before extraction of total RNA or chromatin (see below).

#### Nucleofection in vitro transfection

R22 cells were transfected using the Amaxa™ 4D-Nucleofector™ Optimization Protocol for primary cells (Lonza, Morristown, NJ). R22 cells were grown in culture plates containing DMEM/High glucose supplemented with 10% FBS in a humidified 37’C/5% CO_2_ incubator. Cells were harvested by trypsinization and 6×10^6^ cell were resuspended in 82 ul of 4D-nucleofector solution with 18 ul of supplement solution and 6ug of four different plasmids: empty pcDNA (control), Eed-pcDNA, pKdm6b-pcDNA and pEed-pcDNA/Kdm6b-pcDNA. Eed-pcDNA was described previously by us (8) and Kdm6b-pcDNA was obtained from Addgene (Plasmid # 24167). Each mix was transferred to a Nucleocuvette and electroporated using the 4D-nucleofector Core Unit (LONZA) using the program CA-137. After the electroporation procedure, cells were incubated at room temperature for 5 minutes and then transferred to 10mm culture dishes and incubated for 48 hours in DMEM/High glucose media supplemented with 10% FBS. Transfection efficiency was tested using R22 cells electroporated with pEGFP vector. All transfections where performed at three different times and in triplicate.

### RNA extraction, reverse transcription, and quantitative (q)PCR

Total RNA was extracted from tissues (MBH) and R22 cells using the RNeasy mini kit (Qiagen, Valencia, CA) following the manufacturer’s instructions. RNA concentrations were determined by spectrophotometric trace (Nanodrop, ThermoScientific, Wilmington, DE). Total RNA (2000 ng) was transcribed into cDNA in a volume of 20 μl using 4 U Omniscript reverse transcriptase (Qiagen). To determine the relative abundance of the mRNAs of interest, we used the SYBR GreenER™ qPCR SuperMix system (Invitrogen, Carlsbad, CA). Primers for PCR amplification (**Supplementary Table 5**) were designed using the PrimerSelect tool of DNASTAR 14 software (Madison, WI) or the NCBI online Primer-Blast program. PCR reactions were performed in a total volume of 10 μl containing 1 μl of diluted cDNA or a reference cDNA sample (see below), 5 μl of SYBR GreenER™ qPCR SuperMix and 4 μl of primers mix (1 μM of each gene specific primer). The PCR conditions used were 95°C for 5 min, followed by 40 cycles of 15 sec at 95°C and 60 sec at 60°C. To confirm the formation of a single SYBR Green-labeled PCR amplicon, the PCR reaction was followed by a three-step melting curve analysis consisting of 15 sec at 95°C, 1 min at 60°C, ramping up to 95°C at 0.5°C/sec, detecting every 0.5 sec and finishing for 15 sec at 95°C, as recommended by the manufacturer. All qPCR reactions were performed using a QuantStudio 12K Real-Time PCR system; threshold cycles (CTs) were detected by QuantStudio 12K Flex software. Relative standard curves were constructed from serial dilutions (1/2 to 1/500) of a pool of cDNAs generated by mixing equal amounts of cDNA from each sample. The CTs from each sample were referred to the relative standard curve to estimate the mRNA content/sample; the values obtained were normalized for procedural losses using glyceraldehyde-3-phosphate dehydrogenase (*GAPDH*) mRNA or peptidylprolyl isomerase A (Ppia) as the normalizing unit.

### Massively parallel RNA sequencing (RNA-seq)

Total RNA from ARC-ME fragments derived from rats receiving LV-EED or LV-GFP particles into the ARC and incubated *in vitro* to examine changes in pulsatile GnRH release was subjected to RNA-seq. The RNA-seq procedure was carried out by the OHSU Massively Parallel Sequencing Shared Resource. RNA-seq libraries were prepared using the TruSeq Stranded protocol with ribosomal reduction (Illumina, San Diego, CA). Briefly, 600 ng of total RNA per sample were depleted of ribosomal RNA using RiboZero capture probes (Illumina). The purified RNA was then fragmented using divalent cations and heat, and the fragmented RNA was used as template for reverse transcription using random hexamer primers. The resulting cDNAs were enzymatically treated to blunt the ends, and a single “A” nucleotide was added to the 3’ ends to facilitate adaptor ligation. Standard six-base pair Illumina adaptors were ligated to the cDNAs and the resulting DNA was amplified by 12 rounds of PCR. All of the above procedures were carried out following the protocol provided by Illumina. Unincorporated material was removed using AMPure XP beads (BeckmanCoulter, Brea, CA). Libraries were profiled on a Bioanalyzer instrument (Agilent, Santa Clara, CA) to verify: a) the distribution of DNA sizes in the library, and b) the absence of adapter dimers. Library titers were determined using real time PCR (Kapa Biosystems, Wilmington, MA) on a StepOnePlus Real Time System (ThermoFisher, Waltham, MA). Libraries were mixed to run four samples per lane on the HiSeq 2500 (Illumina). Sequencing was done using a single-read 100-cycle protocol. The resulting base call files (.bcl) were converted to standard fastq formatted sequence files using Bcl2Fastq (Illumina). Sequencing quality was assessed using FastQC (Babraham Bioinformatics, Cambridge, UK). The RNA-seq data is available at NCBI under the accession number GSE102471.

### RNAseq data analysis

To determine the differential expression of genes in LV-GFP and LV-EED containing ARC-ME fragments we used the gene-level edgeR (43) analysis package. We performed an initial trimming and adapter removal pass using Trimmomatic (44). Reads that passed Trimmomatic processing were aligned to the rn6 build of the rat genome with Bowtie2/Tophat2 (45, 46), and assigned to gene-level genomic features with the Rsubread featureCounts package based on the Ensembl 83 annotation set. Differential expression between LV-GFP and LV-EED injected groups was analyzed using the generalized linear modeling approaches implemented in edgeR (13). Lists of differentially expressed genes/transcripts were identified based on significance of pairwise comparisons. A subset of genes found to be differentially expressed was selected for subsequent RT-qPCR confirmation.

### WGCNA Analysis of RNA-seq Data

We used the Weighted-Gene Co-Expression Analysis method for discovery of co-expressed modules of genes in our RNA-seq data. Data were transformed to log2 counts-per-million estimates using the voom function in edgeR. We then determined the 5000 most variable genes across the pooled control and EED-overexpressing samples and applied the WGCNA pipeline to those count estimates. We utilized a signed network and specified a minimum module size of 100 genes; all other parameters were set at defaults. The eigengene summary metric of overall module expression was used to visualize the trend in differential expression of the genes in each module between control and EEDoverexpressing samples. We also performed functional enrichment analysis using the DAVID tool (15, 16) for the genes in each WGCNA module. We utilized the human orthologs of these genes as determined by Ensembl for this analysis due to the superior annotation of the human genome; genes that did not have an assigned ortholog were dropped from the analysis. Overrepresented annotation categories for each set of genes were defined as categories with an FDR-value of 5% or less as reported by DAVID’s modified Fisher Exact Test procedure.

### Open Array-Real Time PCR

We used the OpenArray-qPCR platform to measure changes in relative expression for 224 genes studied after perturbing the system by overexpressing EED in the ARC (rats injected on PND21 and euthanized on PND28; ARC-ME fragments collected after a 4h incubation period to measure changes in pulsatile GnRH release). One thousand ng of total RNA from each ARC-ME fragment were reverse transcribed (RT) using the Omni RT Kit (Qiagen, Valencia, CA) in the presence of random hexamer primers (Invitrogen, Carlsbad, CA), as recommended by the manufacturer. The resulting cDNA was diluted 4 times with H2O and mixed with 2X TaqMan OpenArray Real Time PCR Master Mix (Life Technologies, Grand Island, NY) at a ratio of 3.8:1.2 (PCR mix:cDNA). The mix was loaded into custom made (12X224 probes) OpenArray plates (for target genes probe numbers and lengths of amplicon see (20)) using the Quant Studio OpenArray AccuFill platform and the PCR reactions were performed in a QuantStudio 12K Flex Real-Time PCR System (Applied Biosystems, Foster City, CA).

### Open Array Analysis

Raw data were extracted from the QuantStudio 12K Flex software and analyzed using R. CT values were converted to relative expression levels for further analysis using a standard delta-delta transformation. Genes with unusually high variability and/or multiple missing values were dropped at this stage, resulting in 134 genes. We then performed tests for differential expression using the Student’s unpaired T-test; missing values and outliers were dropped from these analyses. Resultant p-values were adjusted for multiple testing using the Benjamini-Hochberg analysis. For further analyses, we imputed missing expression values and the values of removed outliers using k-nearest neighbor imputation using the impute R package before correction.

### Gene Network Analysis

We utilized a combination of data from the OpenArray platform and standard RT-PCR experiments targeting important genes that were of low quality from the OpenArray runs (e.g. Kiss1) to perform an inference of a network of strong gene co-expression relationships. After correction, strong co-expression networks were derived from the data using a compressive sensing based-method previously described (21); briefly, the method generates all possible one-gene partial correlation matrices from the data (via the ppcor package in R), then utilizes a CLIME-based approach (47) as implemented in the R package clime to approximate inverse matrices for each partial correlation matrix. Networks based on a range of values of the regularization parameter lambda for CLIME are generated; we then select a representative network for each inverse partial correlation network based on highest scale-free fit. Co-expression relationships that are preserved in the 95^th^ percentile or more of the distribution of edge preservation in the inverse partial correlation networks after thresholding for interaction strength are included in the overall network. This process was used to construct a network based on the pooled data from controls and from EED-overexpressing samples. Networks were visualized, analyzed and compared using R and Cytoscape 3.1.1 (www.cytoscape.org).

We also utilized the GeneMANIA (29) gene interaction database to find and visualize known interactions between differentially expressed glutamatergic-related and potassium channel genes located in WGCNA modules of interest, along with the *Kdm6b* second-neighbor genes *Nell2* and *Grm7*. We utilized the human orthologs of these genes for this analysis and the default co-expression, genetic interaction, physical interaction and pathway data sets of the database.

### Chromatin Immunoprecipitation (ChIP) Assay

To assess the recruitment of EED to putative target gene promoters, and the changes in H3K27me3 association to these promoters caused by EED overexpression, we performed ChIP assays using chromatin extracted from R22 immortalized hypothalamic cells overexpressing EED. The ChIP procedure was carried out essentially as previously described (8, 48, 49). The antibodies used (2-5 μg/reaction) are listed in **Supplementary Table 6**.

### qPCR detection of Chromatin Immunoprecipitated DNA

The proximal promoter regions of the genes of interest were amplified by qPCR. **Supplementary Table 5** lists the accession numbers of the genes analyzed, as well as the chromosomal position of the 5’-flanking region amplified, using the position of the transcription start site (TSS) as the reference point. The primer sequences (Eurofins MWG Operon, Huntsville, Al) used to amplify immunoprecipitated DNA fragments are also shown in **Supplementary Table 5**. PCR reactions were performed using 1 μl of each immunoprecipitate (IP) or input samples (see below), primer mix (1 μM each primer), and SYBR Green Power Up Master Mix™ (Thermo Fisher, Waltham, MA) in a final volume of 10 μl. Input samples consisted of 10% of the chromatin volume used for immunoprecipitation. The thermocycling conditions used were: 95°C for 5 min, followed by 40 cycles of 15 sec at 95°C and 60 sec at 60°C. Data are expressed as % of IP signal / Input signal.

### Statistics

All statistical analyses were performed using SigmaStat software (Systat Software Inc., San Jose, CA). The differences between several groups were analyzed by one-way ANOVA followed by the Student-Newman-Keuls multiple comparison test for unequal replications. The Student’s t test was used to compare two groups. When comparing percentages, groups were subjected to arc–sine transformation before statistical analysis to convert them from a binomial to a normal distribution (50). A p value of < 0.05 was considered statistically significant.

## Acknowledgements

This work was supported by grants from the US National Science Foundation (NSF: IOS1121691) to S.R.O, the National Institute of Health (1R01HD084542) to S.R.O and A.L, and 8P51OD011092 for the operation of the Oregon National Primate Research Center. C.A.T and H.W. were supported by NIH Training grants T32-HD007133 and T32 DK 7680. C.A.T was also supported by NRSA 1 F32 HD086904.

## Author Contributions

H.W., S.R.O. and A.L designed the project and wrote the paper, which was revised by the rest of the authors. A.L. conducted and coordinated the molecular and physiological experiments, helped with in the intrahypothalamic injections of lentiviruses and performed the incubation of hypothalamic tissue. S.R.O. coordinated the project, performed the intrahypothalamic injections of lentiviruses, and helped with the incubation of hypothalamic tissue. C.A.T. and C.F.A measured mRNAs by qPCR, performed the ChIP assays and helped in preparing the figures. H.W. performed the *in silico* analysis of microarray and RNA-seq data, the statistical analysis of physiological data and gene expression, and carried out the gene co-expression network analyses.

## Legends to Figures

**Supplementary Figure 1: Changes in gene expression elicited by EED overexpression in hypothalamic R22 cells.** Loss of expression of selected epigenetic, glutamatergic-related and potassium channel genes in hypothalamic R22 cells. Cells were infected with of LV-GFP or LV-EED, isolated by FACS and expanded prior mRNA levels measured by qPCR. Results are expressed as fold change with respect to control values. * = p < 0.05, ** p < 0.01, *** p < 0.001 vs. LV-GFP-Control treated cells (Student’s t-test) (n=4 per group).

**Supplementary Table 1:** DAVID functional analysis of genes in modules from WGCNA of the rat ARC nucleus.

**Supplementary Table 2:** Differential expression analysis of RNAseq data from LV-Control and LV-EED rat ARC nucleus.

**Supplementary Table 3:** Differential expression analysis of custom made OpenArray data from LV-Control and LV-EED rat ARC nucleus.

**Supplementary Table 4:** Partial correlation analysis of ARC nucleus gene expression with GnRH release.

**Supplementary Table 5:** Primers used for OpenArray, RT-qPCR and CHIP-PCR.

**Supplementary Table 6:** Antibodies used for CHIP assays.

## References

1. Ojeda SR, Skinner MK. Puberty in the rat. In: Neill JD, editor. The Physiology of Reproduction (3rd Edition). San Diego: Academic Press/Elsevier; 2006. p. 2061–126.

2. Lomniczi A, Ojeda SR. A role for glial cells of the neuroendocrine brain in the central control of female sexual development. In: Parpura V, Haydon P, editors. Astrocytes in (Patho)Physiology of the Nervous System. Springer, NY 2009. p. 487–511.

3. Ojeda SR, Dubay C, Lomniczi A, Kaidar G, Matagne V, Sandau US, et al. Gene networks and the neuroendocrine regulation of puberty. MolCell Endocrinol. 2010;324(1-2):3–11.

4. Navarro VM, Gottsch ML, Wu M, Garcia-Galiano D, Hobbs SJ, Bosch MA, et al. Regulation of NKB Pathways and Their Roles in the Control of Kiss1 Neurons in the Arcuate Nucleus of the Male Mouse. Endocrinology. 2011;152(11):4265–75.

5. Lehman MN, Coolen LM, Goodman RL. Minireview: kisspeptin/neurokinin B/dynorphin (KNDy) cells of the arcuate nucleus: a central node in the control of gonadotropin-releasing hormone secretion. Endocrinology. 2010;151(8):3479–89.

6. Beale KE, Kinsey-Jones JS, Gardiner JV, Harrison EK, Thompson EL, Hu MH, et al. The physiological role of arcuate kisspeptin neurons in the control of reproductive function in female rats. Endocrinology. 2014;155(3):1091–8.

7. Wakabayashi Y, Nakada T, Murata K, Ohkura S, Mogi K, Navarro VM, et al. Neurokinin B and dynorphin A in kisspeptin neurons of the arcuate nucleus participate in generation of periodic oscillation of neural activity driving pulsatile gonadotropin-releasing hormone secretion in the goat. Journal of Neuroscience. 2010;30(8):3124–32.

8. Lomniczi A, Loche A, Castellano JM, Ronnekleiv OK, Bosh M, Kaidar G, et al. Epigenetic control of female puberty. Nature Neuroscience. 2013;16:281–9.

9. Schwartz YB, Pirrotta V. Polycomb silencing mechanisms and the management of genomic programmes. NatRevGenet. 2007;8(1):9–22.

10. Kohler C, Villar CB. Programming of gene expression by Polycomb group proteins. Trends Cell Biol. 2008;18(5):236–43.

11. Simon JA, Kingston RE. Mechanisms of polycomb gene silencing: knowns and unknowns. NatRevMolCell Biol. 2009;10(10):697–708.

12. Langfelder P, Horvath S. WGCNA: an R package for weighted correlation network analysis. BMCBioinformatics. 2008;9:559.

13. McCarthy DJ, Chen Y, Smyth GK. Differential expression analysis of multifactor RNA-Seq experiments with respect to biological variation. Nucleic Acids Research. 2012;40(10):4288–97.

14. R: A Language and Environment for Statistical Computing. R Foundation for Statistical Computing. Viena, Austria 2017.

15. Huang dW, Sherman BT, Lempicki RA. Bioinformatics enrichment tools: paths toward the comprehensive functional analysis of large gene lists. Nucleic Acids Research. 2009;37(1):1–13.

16. Huang dW, Sherman BT, Lempicki RA. Systematic and integrative analysis of large gene lists using DAVID bioinformatics resources. NatProtoc. 2009;4(1):44–57.

17. Wan J, Zhan J, Li S, Ma J, Xu W, Liu C, et al. PCAF-primed EZH2 acetylation regulates its stability and promotes lung adenocarcinoma progression. Nucleic Acids Research. 2015;43(7):3591–604.

18. Hendrich B, Bird A. Identification and characterization of a family of mammalian methyl-CpG binding proteins. MolCell Biol. 1998;18(11):6538–47.

19. Schwartz YB, Pirrotta V. A new world of Polycombs: unexpected partnerships and emerging functions. NatRevGenet. 2013;14(12):853–64.

20. Nottke A, Colaiacovo MP, Shi Y. Developmental roles of the histone lysine demethylases. Development. 2009;136(6):879–89.

21. Lomniczi A, Wright H, Castellano JM, Matagne V, Toro CA, Ramaswamy S, et al. Epigenetic regulation of puberty via Zinc finger protein-mediated transcriptional repression. NatCommun. 2015;6:10195.

22. Ha CM, Choi J, Choi EJ, Costa ME, Lee BJ, Ojeda SR. NELL2, a neuron-specific EGF-like protein, is selectively expressed in glutamatergic neurons and contributes to the glutamatergic control of GnRH neurons at puberty. Neuroendocrinology. 2008;88(3):199–211.

23. Rice JC, Briggs SD, Ueberheide B, Barber CM, Shabanowitz J, Hunt DF, et al. Histone methyltransferases direct different degrees of methylation to define distinct chromatin domains. MolCell. 2003;12(6):1591–8.

24. Schultz DC, Ayyanathan K, Negorev D, Maul GG, Rauscher FJ, III. SETDB1: a novel KAP-1-associated histone H3, lysine 9-specific methyltransferase that contributes to HP1-mediated silencing of euchromatic genes by KRAB zinc-finger proteins. Genes and Development. 2002;16(8):919–32.

25. De Sousa RT, Loch AA, Carvalho AF, Brunoni AR, Haddad MR, Henter ID, et al. Genetic Studies on the Tripartite Glutamate Synapse in the Pathophysiology and Therapeutics of Mood Disorders. Neuropsychopharmacology. 2017;42(4):787–800.

26. Goldstein SA, Bayliss DA, Kim D, Lesage F, Plant LD, Rajan S. International Union of Pharmacology. LV. Nomenclature and molecular relationships of two-P potassium channels. PharmacolRev. 2005;57(4):527–40.

27. Ramaswami M, Gautam M, Kamb A, Rudy B, Tanouye MA, Mathew MK. Human potassium channel genes: Molecular cloning and functional expression. MolCell Neurosci. 1990;1(3):214–23.

28. Boettger MK, Till S, Chen MX, Anand U, Otto WR, Plumpton C, et al. Calcium-activated potassium channel SK1-and IK1-like immunoreactivity in injured human sensory neurones and its regulation by neurotrophic factors. Brain. 2002;125(Pt 2):252–63.

29. Warde-Farley D, Donaldson SL, Comes O, Zuberi K, Badrawi R, Chao P, et al. The GeneMANIA prediction server: biological network integration for gene prioritization and predicting gene function. Nucleic Acids Research. 2010;38(Web Server issue):W214–W20.

30. Maeda K, Ohkura S, Uenoyama Y, Wakabayashi Y, Oka Y, Tsukamura H, et al. Neurobiological mechanisms underlying GnRH pulse generation by the hypothalamus. Brain Res. 2010;1364:103–15.

31. Moenter SM, DeFazio AR, Pitts GR, Nunemaker CS. Mechanisms underlying episodic gonadotropin-releasing hormone secretion. Front Neuroendocrinol. 2003;24(2):79–93.

32. Conte FA, Grumbach MM, Kaplan SL, Reiter EO. Correlation of luteinizing hormone-releasing factor-induced luteinizing hormone and follicle-stimulating hormone release from infancy to 19 years with the changing pattern of gonadotropin secretion in agonadal patients: relation to the restraint of puberty. J Clin Endocrinol Metab. 1980;50(1):163–8.

33. Waldhauser F, Weissenbacher G, Frisch H, Pollak A. Pulsatile secretion of gonadotropins in early infancy. Eur J Pediatr. 1981;137(1):71–4.

34. Rasmussen DD. Episodic gonadotropin-releasing hormone release from the rat isolated median eminence in vitro. Neuroendocrinology. 1993;58(5):511–8.

35. Wilson RC, Kesner JS, Kaufman JM, Uemura T, Akema T, Knobil E. Central electrophysiologic correlates of pulsatile luteinizing hormone secretion in the rhesus monkey. Neuroendocrinology. 1984;39(3):256–60.

36. Thiery JC, Pelletier J. Multiunit activity in the anterior median eminence and adjacent areas of the hypothalamus of the ewe in relation to LH secretion. Neuroendocrinology. 1981;32(4):217–24.

37. Navarro VM. Interactions between kisspeptins and neurokinin B 1. AdvExpMedBiol. 2013;784:325–47.

38. Navarro VM, Gottsch ML, Chavkin C, Okamura H, Clifton DK, Steiner RA. Regulation of gonadotropin-releasing hormone secretion by kisspeptin/dynorphin/neurokinin B neurons in the arcuate nucleus of the mouse. Journal of Neuroscience. 2009;29(38):11859–66.

39. Roth CL, Mastronardi C, Lomniczi A, Wright H, Cabrera R, Mungenast AE, et al. Expression of a tumor-related gene network increases in the mammalian hypothalamus at the time of female puberty. Endocrinology. 2007;148(11):5147–61.

40. Cao R, Wang L, Wang H, Xia L, Erdjument-Bromage H, Tempst P, et al. Role of histone H3 lysine 27 methylation in polycomb-group silencing. Science. 2002;298:1039–43.

41. Saha B, Home P, Ray S, Larson M, Paul A, Rajendran G, et al. EED and KDM6B coordinate the first mammalian cell lineage commitment to ensure embryo implantation. MolCell Biol. 2013;33(14):2691–705.

42. Chung N, Bogliotti YS, Ding W, Vilarino M, Takahashi K, Chitwood JL, et al. Active H3K27me3 demethylation by KDM6B is required for normal development of bovine preimplantation embryos. Epigenetics. 2017;12(12):1048–56.

43. Robinson MD, McCarthy DJ, Smyth GK. edgeR: a Bioconductor package for differential expression analysis of digital gene expression data. Bioinformatics. 2010;26(1):139–40.

44. Bolger AM, Lohse M, Usadel B. Trimmomatic: a flexible trimmer for Illumina sequence data. Bioinformatics. 2014;30(15):2114–20.

45. Langmead B, Salzberg SL. Fast gapped-read alignment with Bowtie 2. NatMethods. 2012;9(4):357–9.

46. Kim D, Pertea G, Trapnell C, Pimentel H, Kelley R, Salzberg SL. TopHat2: accurate alignment of transcriptomes in the presence of insertions, deletions and gene fusions. Genome Biol. 2013;14(4):R36.

47. Cai TT, Liu W, Luo X. A constraind L1 minimization approach for sparse precision matrix estimation. J Amer Stat Assoc. 2011;106(494):594–607.

48. Mueller JK, Dietzel A, Lomniczi A, Loche A, Tefs K, Kiess W, et al. Transcriptional regulation of the human KiSS1 gene. Molecular and Cellular Endocrinology. 2011;342(1-2):8–19.

49. Mueller JK, Koch I, Lomniczi A, Loche A, Rulfs T, Castellano JM, et al. Transcription of the human EAP1 gene is regulated by upstream components of a puberty-controlling Tumor Suppressor Gene network. MolCell Endocrinol. 2012;351(2):184–98.

50. Zar JH. Biostatistical Analysis, 2nd Edition. Englewood Cliffs, NJ: Prentice Hall; 1984.

