## Supplementary figures and images for "Polycomb represses a gene network controlling puberty via modulation of histone demethylase *Kdm6b* expression"

### Suppl. Fig. 1 mRNA R22-0 v3 copy.jpg

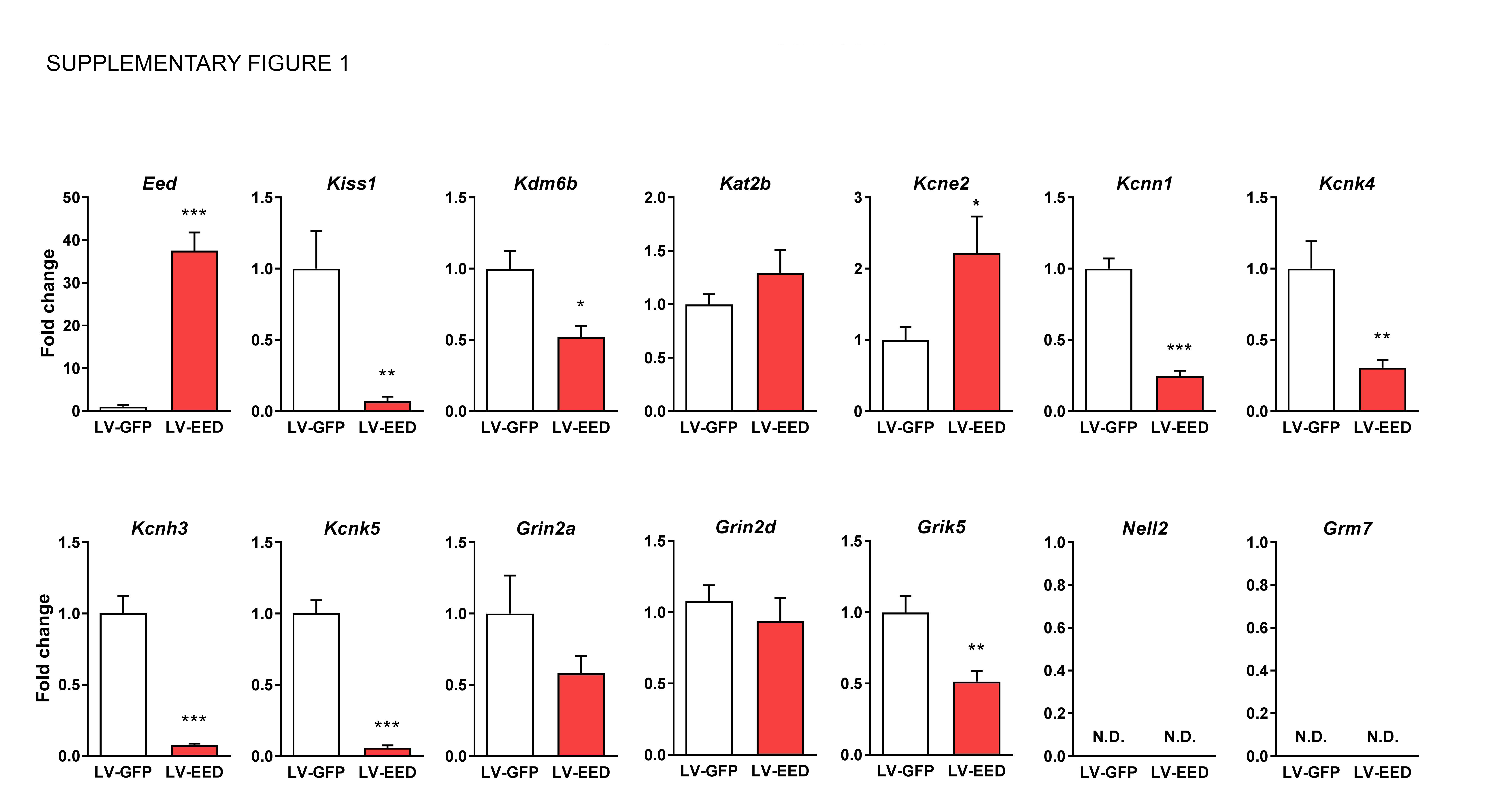
